# The prognostic potential of alternative transcript isoforms across human tumors

**DOI:** 10.1101/036947

**Authors:** Juan L. Trincado, E. Sebestyén, A. Pagés, E. Eyras

**Affiliations:** Universitat Pompeu Fabra (UPF), Dr. Aiguader 88, E08003 Barcelona, Spain; The Firc Institute of Molecular Oncology (IFOM), Via Adamello 16, 20139 Milan, Italy; Catalan Institution for Research and Advanced Studies (ICREA), Passeig Lluís Companys 23, E08010 Barcelona, Spain

## Abstract

**Background:** Phenotypic changes during cancer progression are associated to alterations in gene expression, which can be exploited to build molecular signatures for tumor stage identification and prognosis. However, it is not yet known whether the relative abundance of transcript isoforms may be informative for clinical stage and survival.

**Methods:** Using information theory and machine learning methods, we integrated RNA sequencing and clinical data from The Cancer Genome Atlas project to perform the first systematic analysis of the prognostic potential of transcript isoforms in 12 solid tumors to build new predictive signatures for stage and prognosis. This study was also performed in breast tumors according to estrogen receptor status and melanoma tumors with proliferative and invasive phenotypes.

**Results:** Transcript isoform signatures accurately separate early from late stage and metastatic from non-metastatic tumors, and are predictive of the survival of patients with undetermined lymph node invasion or metastatic status. These signatures show similar, and sometimes better, accuracies compared with known gene expression signatures, and are largely independent of gene expression changes. Furthermore, we show frequent transcript isoform changes in breast tumors according to estrogen receptor status, and in melanoma tumors according to the invasive or proliferative phenotype, and derive accurate predictive models of stage and survival within each patient subgroup.

**Conclusions:** Our analyses reveal new signatures based on transcript isoform abundances that characterize tumor phenotypes and their progression independently of gene expression. Transcript isoform signatures appear especially relevant to determine lymph node invasion and metastasis, and may potentially contribute towards current strategies of precision cancer medicine.

## Introduction

Tumors advance through stages that are generally characterized by their size and spread to lymph nodes and other parts of the body [1]. Establishing the stage of a tumor is critical to determine patient prognosis and to select the appropriate therapeutic strategy [2]. Even though stage is generally defined from a number of tests carried out on a patient, this information may sometimes be incomplete or inconclusive. Advances in the molecular characterization of tumors have lead to improvements in stage classification and clinical management of patients [3]. Although tumors originate primarily from genetic lesions, their progression involves other molecular transformations, which are related to the activation of specific aggressive phenotypes, like tumor spread and metastasis, and are often reflected in gene expression changes [4,5]. Accordingly, the development of gene expression signatures has been instrumental to complement and improve stage identification and prognosis [6–9]. On the other hand, gene expression summarizes the output of RNA transcripts from a gene locus, which is mostly explained by one transcript isoform [10]. Furthermore, we described before how solid tumors present frequent changes in the relative abundances of isoforms in comparison to normal tissues [11]. This prompts the question of whether transcript isoform changes, which remain largely unexplored as predictive signatures of tumor stage and survival, could hold relevant novel mechanisms of tumor progression. We investigated the predictive potential of the relative abundances of transcript isoforms for tumor staging and clinical outcome in 12 different tumor types, integrating RNA sequencing and clinical annotation data for 12 tumor types from The Cancer Genome Atlas (TCGA) project. Our analyses revealed new signatures that characterize tumor phenotypes and their progression largely independent of gene expression. Knowledge about the relative abundance of transcript isoforms in tumors can potentially help predicting stage and clinical outcome and contribute towards current molecular strategies in precision cancer medicine.

## Results

### Relative abundances of transcript isoforms are predictive of tumor stage

We considered the standard clinical annotation for tumors based on the tumor size (T), lymph-node involvement (N) metastatic status (M) and combined stage (S), for 4339 patient samples from 12 different tumor types from TCGA (Additional file 1). For each tumor type, we considered the comparison of the transcriptomes between groups of samples in early and late stage groups according to each stage class independently. That is, for metastasis, we compared non-metastatic samples (M0) against metastatic ones (M1), whereas for the tumor size (T), lymph-node involvement (N) and stage (S) annotations, we compared early and late stages (groups described in Table 1) (Methods). We first calculated the set of transcripts whose relative abundance, measured as percent spliced in (PSI) values, present the best discriminant potential between these groups by using information-based measures with a subsampling strategy to ensure balanced comparisons, and selecting (Fig. 1a) (Fig. S1a in Additional file 2). Additionally, we considered only those transcripts that on average change PSI more than 10% between groups, i.e. |ΔPSI| > 0 (Methods). These produced a variable number of transcript isoforms per tumor type and clinical annotation that discriminate between early and late stages, or between M0 and M1 (Additional file 3).

**Table 1.** Number of samples analyzed for each tumor type and stage.

| Tumor type | Acronym | T |  | N |  | M |  | S |  |
| --- | --- | --- | --- | --- | --- | --- | --- | --- | --- |
|  |  | Early | Late | Early | Late | Early | Late | Early | Late |
| Breast invasive carcinoma | BRCA | 256<br>(T1) | 147<br>(T3,T4) | 455<br>(N0) | 171<br>(N2,N3) | 836<br>(M0) | 15<br>(M1) | 164<br>(S1) | 15<br>(S4) |
| Colon adenocarcinoma | COAD | 45<br>(T1,T2) | 31<br>(T4) | 149<br>(N0) | 39<br>(N2) | 179<br>(M0) | 33<br>(M1) | 40<br>(S1) | 34<br>(S4) |
| Head and neck squamous cell carcinoma | HNSC | 35<br>(T1) | 110<br>(T4) | 166<br>(N0) | 166<br>(N2,N3) |  |  | 77<br>(S1,S2) | 169<br>(S4) |
| Kidney chromophobe | KICH | 20<br>(T1) | 19<br>(T3,T4) |  |  |  |  | 20<br>(S1) | 19<br>(S3,S4) |
| Kidney renal clear cell carcinoma | KIRC | 245<br>(T1) | 186<br>(T3,T4) | 233<br>(N0) | 16<br>(N1) | 419<br>(M0) | 77<br>(M1) | 240<br>(S1) | 78<br>(S4) |
| Kidney renal papillary carcinoma | KIRP | 71<br>(T1) | 38<br>(T3,T4) | 23<br>(N0) | 16<br>(N1,N2) |  |  | 66<br>(S1) | 38<br>(S3,S4) |
| Lung squamous cell carcinoma | LUSC | 93<br>(T1) | 59<br>(T3,T4) | 242<br>(N0) | 37<br>(N2,N3) |  |  | 195<br>(S1) | 76<br>(S3,S4) |
| Lung adenocarcinoma | LUAD | 137<br>(T1) | 57<br>(T3,T4) | 281<br>(N0) | 70<br>(N2,N3) | 307<br>(M0) | 22<br>(M1) | 99<br>(S1) | 242<br>(S3,S4) |
| Ovarian serous cystadenocarcinoma | OV |  |  |  |  |  |  | 18<br>(S2) | 243<br>(S4) |
| Prostate adenocarcinoma | PRAD | 69<br>(T2) | 93<br>(T3,T4) | 129<br>(N0) | 14<br>(N1) |  |  |  |  |
| Skin cutaneous melanoma | SKCM |  |  |  |  | 68<br>(M0) | 17<br>(M1) |  |  |
| Thyroid Carcinoma | THCA | 137<br>(T1) | 179<br>(T3,T4) | 220<br>(N0) | 211<br>(N1) |  |  | 270<br>(S1) | 48<br>(S4) |

The number of samples used for the comparison early vs late are indicated for each annotation T, N, M, S. Stages I, II, III and IV are indicated as S1, S2, S3 and S4. Comparisons were performed between the earliest and latest available stage groups, with some exceptions for which adjacent stages were added to have enough samples for comparison. Empty cells correspond to cases not tested due to lack of sufficient samples or complete lack of annotation in the samples.

**Figure 1.**
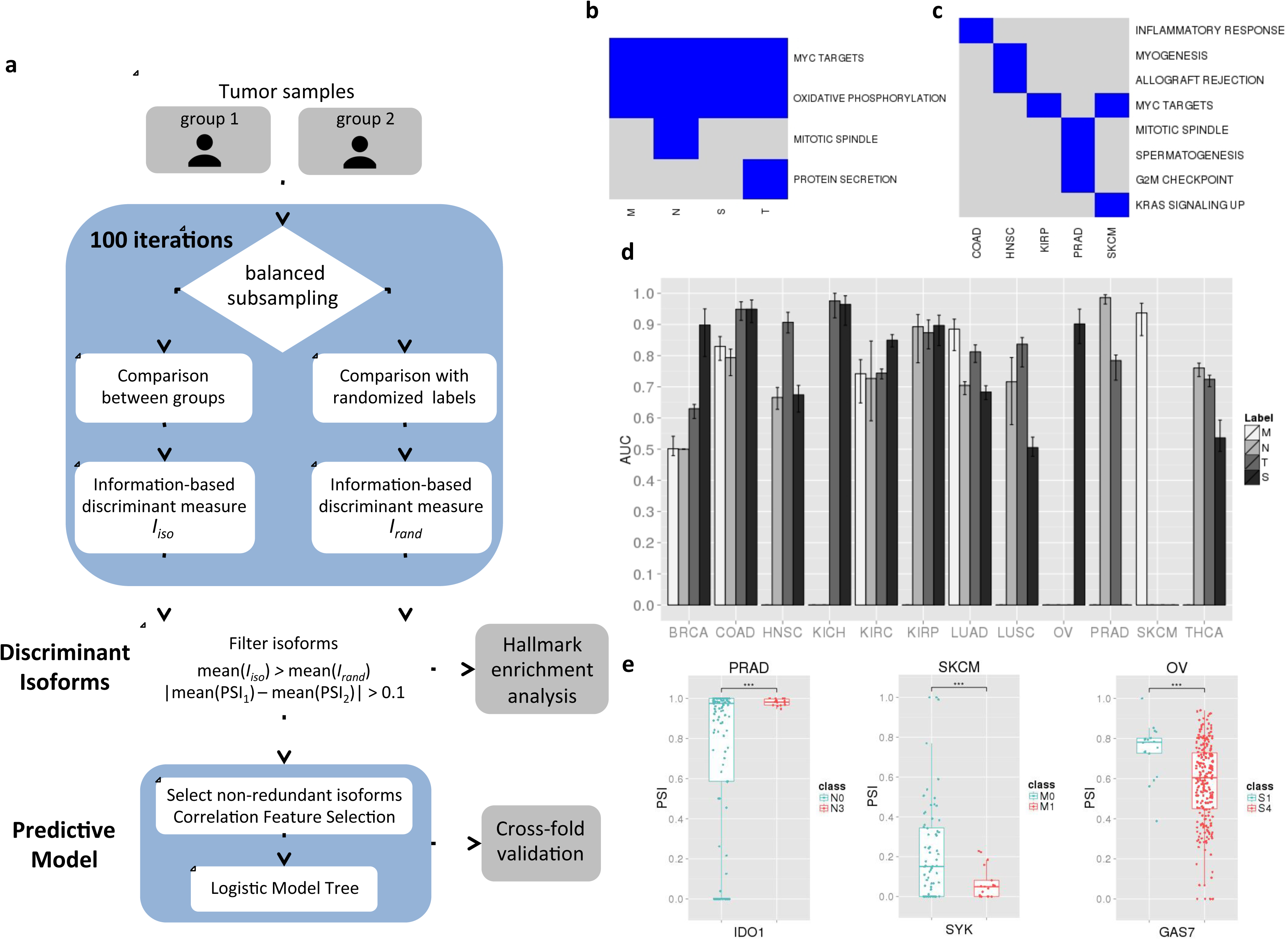
**(a)** Workflow to obtain discriminant transcript isoforms and predictive models. Given two patient groups, we subsampled two equal sized subsets, one from each group (e.g. metastatic and non-metastatic), which were compared using information-based measures, denoted as *I*_*iso*_. At each iteration step, the group labels were randomized to obtain an expected measure, denoted as *I*_*rand*_ After 100 iterations, two distributions were produced for each isoform corresponding to observed (*I*_*iso*_) and expected (*I*_*rand*_) values. Transcript isoforms with a difference of mean PSI values >0.1 in absolute value between the two patient groups and with a positive difference of the means of the observed and expected distributions for all information-based measures used were then considered as discriminant, which were then used to evaluate enriched cancer hallmarks. Discriminant isoforms were further filtered for redundancy with a Correlation Feature Selection strategy to build a predictive model, which was evaluated using cross-fold validation (Methods). **(b)** Enriched hallmarks in the set of discriminant isoforms for each stage class, metastasis (M), tumor size (T), lymph-node involvement (N) and overall staging (S), using all isoforms selected across all tumor types. **(c)** Enriched hallmarks for each tumor type using all discriminant isoforms selected across all stage classes in each tumor type independently. **(d)** Accuracies of the classifiers for each tumor type for the T, N, M and S annotation, given as the distributions of the areas under the receiving operating characteristic (ROC) curves (AUC). The variation on each bar indicates the minimum and maximum AUC values. Some models are absent due to lack of sufficient samples (Table 1). **(e)** PSI distributions for the transcript isoforms of *IDO1* in PRAD, *SYK* in SKCM and *GAS7* in OV, for the N, M and S models, respectively (Wilcoxon test p-values < 0.001).

To characterize the functional involvement of the found discriminant isoforms, we performed an enrichment analysis of cancer hallmarks (Methods) (Additional file 4). Testing discriminant isoforms for each stage class and tumor type independently yielded frequent enrichment of MYC targets, oxidative phosphorylation, mTORC signaling, DNA repair and Interferon response (Fig. S1b in Additional file 2). Notably, aggregating all tumor types for each clinical class, the discriminant transcripts show enrichment in MYC targets and genes involved in oxidative phosphorylation (Fig. 1b). On the other hand, combining discriminant isoforms from different clinical classes in the same tumor type, only 5 of the 12 tumor types tested show enriched hallmarks (Fig. 1c), which include the enrichment of MYC targets in skin cutaneous melanoma (SKCM) and kidney papillary carcinoma (KIRP). These results indicate that there are frequent transcripts isoform changes in cancer-relevant pathways during tumor progression, many of which may be driven by MYC activity. To test some of our findings, we compared the ΔPSI values of the discriminant transcripts for metastasis in SKCM with the ΔPSI values measured between metastatic (SKMel147) [12] and nonmetastatic (Mel505) [13] melanoma cells (Methods). Of the 958 discriminant isoforms in SKCM, 817 had expression in the cell lines. From these, 504 (61.7%) show a change in PSI, in the same direction and 253 of them have |ΔPSI| > 0.1 in both comparisons (Fig. S1c in Additional file 2).

To build signatures of tumor stage based on transcript isoforms, we applied a multivariate feature selection method on the discriminant isoforms selected before to obtain a nonredundant subset of predictive transcripts, which we used to build Logistic Model Trees (LMT) for each tumor type and stage class (Fig. 1a) (models given in Additional file 5). Each one of these models represents a transcript signature for each stage class and each tumor type. Using cross-validation, the mean accuracy of the models in terms of the area under the ROC curve (AUC) is 0.783 (Fig. 1d), with similar average precision-recall values (Fig. S1d in Additional file 2). T-models show the best accuracies (mean AUC = 0.824), with the models for KIRP, kidney chromophobe (KICH), colon adenocarcinoma (COAD) and neck squamous cell carcinoma (HNSC) being the most accurate (mean AUC > 0.87).

KIRP T-model includes an isoform for *PAX6*. Increased inclusion of exon 5 of this gene has been related to neuronal differentiation [14], which we see associated to late T stage (Fig. S2a in Additional file 2). The best N-models correspond to KIRP and prostate adenocarcinoma (PRAD) (mean AUC > 0.89). KIRP N-model includes an isoform in the MAP kinase MKNK1 (Fig. S2a in Additional file 2), suggesting a similar involvement in cancer as *MKNK2* [15]. PRAD N-model (mean AUC = 0.986) includes an isoform of *IDO1* (Fig. 1e), a gene related to anti tumor defense [16]. The best M-model corresponds to SKCM (mean AUC = 0.93) and includes an isoform change in the transmembrane gene *TM6SF1* (Fig. S2a in Additional file 2) and the tyrosine kinase *SYK* (Fig. 1e). In metastatic melanoma samples, *SYK* shows an increase in the abundance of the long form and a decrease of the short form, as previously observed in breast tumors [17]. Finally, the best S-models correspond to COAD, BRCA, KICH and Ovarian serous cystadenocarcinoma (OV) (AUC > 0.9). Interestingly, OV S-model includes an isoform in the cancer driver *GAS7* (Fig. 1e). In general, we found no overlap between the different stage models. A notable exception is an isoform of *NSUN7* that appears in all models for KIRC with high PSI values at late-stage and an isoform of *SKA3* that appears in the N, T and S models for KIRP, with low PSI values at late stages. The low general overlap is consistent with pathological transformations being associated with multiple molecular alterations.

### Transcript isoform changes are predictive of patient survival

We hypothesized that if the derived transcript signatures provide clinically relevant information, we should find worse clinical outcomes for patients predicted to be at late stage. We thus performed a blind test on those samples that lacked stage annotation, and therefore were not used for building the models, to predict the tumor stage using the model for the corresponding tumor type (Fig. 2a) (Additional file 1). Additionally, we only performed the blind test in those tumor types for which late clinical stage was significantly associated to worse prognosis in the labeled samples (Table 2). There were 40 samples from COAD, 116 from lung adenocarcinoma (LUAD) and 80 from BRCA that lacked M annotation. After prediction with the M-model from each tumor type, we obtained a total of 226 patients predicted as M0 and 10 patients predicted as M1. Aggregating patients according to the predicted metastatic class yielded a significant difference in survival between the two groups (p-value = 0.0079) (Fig. 2c). Regarding lymph node invasion, there were 1 sample from COAD, 10 from LUAD, 82 from KIRP, 247 from KIRC and 74 from HNSC without N annotation. After predicting with the N-models from the corresponding tumor types, 356 and 58 patients were predicted as early and late N, respectively. Survival analysis with the aggregated patients yielded a significant difference between the two predicted groups (p-value = 0.013) (Fig. 2d). Finally, for the S stage, we predicted on a set of 91 samples without S annotation (8 from COAD, 18 from BRCA, 47 from HNSC, 11 from KIRP, 4 from LUSC, 2 from THCA, and 1 from LUAD). This resulted in 47 and 44 samples predicted as early and late, respectively, which showed no difference in survival (p-value = 0.479). These results represent an independent validation of our transcript signatures and provide evidence that the relative abundances of transcripts can be predictive of tumor staging and prognosis.

**Table 2.** Survival analysis between early and late stage patient groups.

| **Tumor type** | **T** | **N** | **M** | **S** |
| --- | --- | --- | --- | --- |
| BRCA | P=0.375 | P=0.00012 | P=0.008 | P=0.0007 |
| COAD | P=0.0011 | P=0.011 | P=1.48e-05 | P=0.012 |
| HNSC | P=0.051 | P=0.0137 |  | P=2.49e-07 |
| KICH | P=0.00896 |  |  | P=0.00896 |
| KIRC | P=2e-15 | P=0.0125 | P=0 | P=0 |
| KIRP | P=0.0043 | P=0.005 |  | P=8.86e-007 |
| LUSC | P=0.029 | P=0.071 |  | P=0.025 |
| LUAD | P=7.02e-09 | P=3.26e-06 | P=0.165 | P=7.02e-09 |
| OV |  |  |  | P=0.0537 |
| PRAD | P=0.456 | P=1 |  |  |
| SKCM |  |  | P=0.418 |  |
| THCA | P=0.324 | P=0.597 |  | P=2.49e-07 |

P-values from the survival test comparing the patient subsets from Table 1. The p-values were obtained using a Cox proportional hazards regression model. Light gray cells indicate comparisons with no significant difference in patient survival. Empty cells correspond to cases not tested due to lack of sufficient samples (see Table 1).

**Figure 2.**
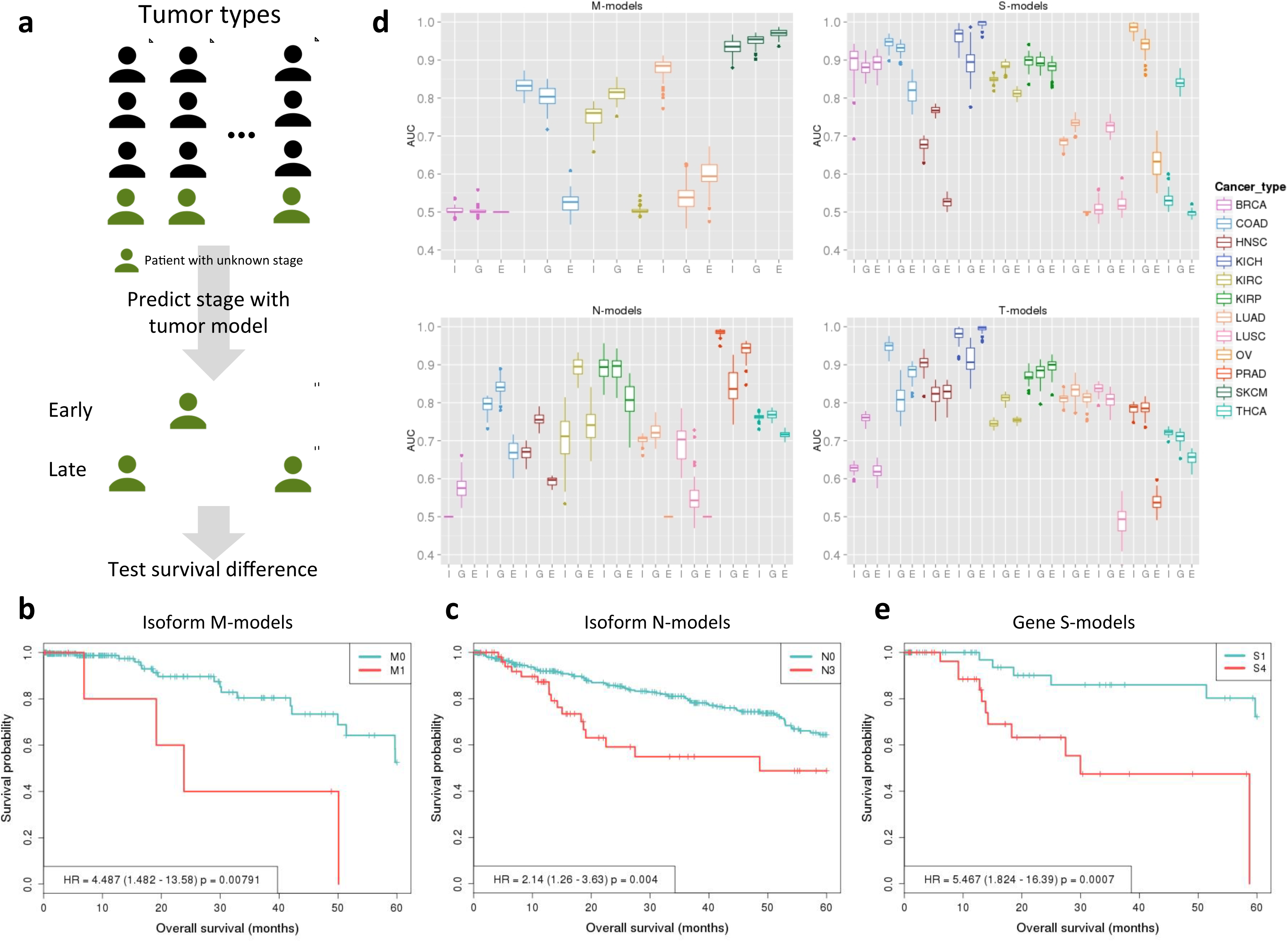
**(a)** Illustration of the blind test on unlabeled patients. Patients without annotated stage were predicted using the model of the corresponding tumor type, for each of the stage classes independently. Patients predicted as early or late were collected into two separate groups and tested for differences in survival. This test was performed for each stage class independently and only using tumor types that showed an association between stage and survival in the labeled patients (Table 2). Figures **(b)** and **(c)** show survival (Kaplan-Meyer) plots associated to the test for M- and N-models, respectively. They indicate the survival percentage (y axis) versus survival in months (x axis) based on the predicted stage on the unannotated samples using the classifier for each corresponding tumor type. The p-value in each plot corresponds to the Cox regression between the two groups and HR indicates the hazards ratio. **(d)** Accuracies of the transcript isoform models (T) compared to the gene (G) and event (E) models. Accuracies are given as boxplots for the distribution of AUC values (y axis) from a 10-fold cross-validation for each tumor type (x axis) for the M, S, N and T models. Tumors for which stage data was missing are not shown (Table 1). **(e)** Survival (Kaplan-Meyer) plot of the early and late stage predictions performed with the gene-based S-models on unannotated samples. The p-value corresponds to the Cox regression between the two groups and HR indicates the hazards ratio.

### No relation of isoform signatures with stromal and immune cell content

To assess whether the purity of the samples could a potential confounding factor of the derived signatures, we tested the correlation between the transcript PSI values of our predictive models against signatures of stromal and immune cell content [18] (Methods). Overall, all signatures showed low correlation with stromal content (mean Pearson |R| < 0.4, Pearson), and all except the N-model in BRCA (Pearson R=0.433) had mean |R|<0.4 with immune cell content (Additional file 6). From the 547 transcript isoforms tested, 95% show a correlation |R|<0.4 (Pearson) for both stromal and immune scores. Among the few cases with |R|>0.5 there is an isoform of *ENAH* (Fig. S2d in Additional file 2), which is present in the T-models in KIRP and COAD and that was previously linked to an invasive phenotype [19]. Recent analyses have shown that clinical stage does not correlated with tumor purity in the TCGA samples [20]. Our analysis further supports those results and indicates that isoform-based signatures of stage do not reflect stromal or immune cell content.

### No universal transcript isoform signature for tumor staging

Our results prompt the question of whether there might be a universal signature of stage and survival based on transcript isoform changes. To test this, we grouped all annotated samples from the different tumor types according to the stage class and applied the same analyses as before. We could only build M and S models due to the lack of common isoforms with discriminant power for the other classes (Additional file 7). The average AUC values for M and S models were lower than before, with mean AUC of 0.5 and 0.685, respectively. Aggregating samples from BRCA, COAD and LUAD, we observed a slight increase in accuracy (mean AUC = 0.702). Similarly, analyzing KIRC, KIRP and KICH samples together, the S-model achieves mean AUC = 0.809. In this case, approximately half of the isoforms were present in the previous models. Finally, analyzing the squamous tumors together (HNSC and LUSC), we derived N and S models with mean AUC = 0.72. For other combinations, we could not find accuracies greater than AUC = 0.5. This indicates that despite some overlapping features across tumor types, there is no common signature for all the tested tumor types.

### Transcript signatures provide better predictions than event-based signatures

We tested whether local alternative splicing events, as opposed to transcript isoform changes, could also be predictive of stage. We applied the same analysis pipeline using PSI values for all events in the same tumor samples. For most of the stage classes we observed similar or smaller accuracy values for events compared to transcript models (average AUC 0.617 vs. 0.778, respectively) (Fig. 2d) (Additional file 8). Only 23.5% of the isoforms in models overlap with at least one alternative splicing event from the event-based models: 16.51% overlap with alternative 5’/3’ splice-sites, mutually exclusive exons, retained introns or cassette exon events, and 6.54% overlap with alternative first or last exon events. Moreover, 82.39% of isoforms in models overlap with at least one of the pre-calculated alternative splicing event. This indicates that a considerable number of changes in exon-intron structures described by the isoform models that are predictive of tumor stage cannot be captured in terms of simple alternative splicing events.

### Transcript signatures provide relevant information about tumor metastasis and lymph node invasion independently of gene expression

Previously proposed molecular classifiers of stage were based on gene expression [7, 8]. We thus tested the relation of our transcript signatures with gene expression. We observed that the proportion of genes with differential expression vary markedly between transcript signatures (Additional file 5) (Methods). For M-models, 9 (18%) genes in the SKCM and 4 (18%) genes in KIRC showed differential expression. For N-models, we only found 3 (14%) in PRAD and 13 (68%) in THCA. In contrast, T-models presented frequent changes across the different tumor types, with 17 (46%) in KIRP, 7 (27%) in to KIRC, 6 (33%) in LUAD, 4 (50%) in THCA and 1 in HNSC (5%). Similarly, S-models also showed frequent DE: 16 (52%) in KIRC, 12 (46%) in KIRP, 1 (25%) in LUAD, and 1 (7%) in BRCA.

Next, we compared the predictive power of transcript and gene expression signatures. We thus applied our pipeline to gene expression values to derive gene-based signatures of stage (Methods). The overall accuracy for gene-based signatures was similar to isoform-based models (average AUC values 0.783 and 0.781 for isoforms and genes, respectively) (Fig. 2d) (Additional file 9). Interestingly, isoforms had better mean accuracies for the M-model in LUAD (0.883 vs 0.535) (Fig. 2d upper left panel) and for the N-model in PRAD (0.986 vs 0.839) (Fig. 2d lower left panel), compared to gene models. In contrast, the gene-based S-model for THCA showed higher accuracy (0.529 vs 0.836) (Fig. 2d upper right panel). Gene and isoform based models generally involved different genes with only few exceptions, including *CD72* in SKCM M-models, *PTGS2* and *VIPR1* in the THCA T-models, *SLC14A1* in COAD S-models, and *DNASE1L3* KICH S-models. Interestingly, gene-based S-models were predictive of survival for samples lacking stage S annotation (p-value = 0.0024) (Fig. 2e), whereas no significant difference in survival was found with the gene-based M and N models (p-values = 0.983 and 0.161, respectively).

The results described above suggest that genes and transcripts provide independent information and may yield better predictors when combined together. We thus built mixed models of gene expression and transcript relative abundance. We started with all gene and transcript discriminant features and selected a non-redundant set of features to build logistic-model trees. The accuracy of these mixed models was on average better (mean AUC = 0.831) than using only transcripts or genes (Fig. S3a in Additional file 2). Notably, the transcript signatures performed better than the mixed signatures for the LUAD M-model (AUC = 0.883 vs 0.814) or the PRAD N-model (AUC = 0.986 vs 0.938). In contrast, the mixed model performed better in the COAD M-model (AUC = 0.831 vs 0.864) and the HNSC S-model (AUC = 0.676 vs 0.778). Additionally, mixed models are able to predict survival differences between early and late stages for the N and S stage classes (p-value = 0.041 and 0.033, respectively) (Figs. S3b and S3c in Additional file 3).

Finally, we compared our transcript signatures with an expression signature of 44 genes built to differentiate metastatic and late stage samples in colon cancer [21] (Methods). The mean AUC values obtained for the metastatic annotation (M) and the overall stage (S) were 0.612 and 0.649, respectively, for the gene expression signature, and 0.82 and 0.94, for our transcript signatures. Notably, none of the genes involved in our transcript models for COAD presented differential expression. Our analyses indicate that changes in the relative abundance of transcripts hold relevant information about tumor transformation independently of gene expression changes.

### Transcript relative abundances as prognostic markers in ER-negative breast tumors

Molecular subtypes in cancer have implications for prognosis and therapy that go beyond the staging system [6, 22, 23]. In breast cancer, tumors that are negative for the Estrogen receptor (ER) have generally worse prognosis, and gene expression signatures are generally less accurate for ER negative than for ER positive tumors [3,7]. To test whether transcript-based signatures could be relevant for ER negative tumors, we separated the samples according to the expression ranking of the ER gene *(ESR1)* into the top (ER+) and bottom (ER-) and 25% (237 samples each) (Fig. 3a). Interestingly, applying our pipeline we identified 2591 discriminant transcript isoforms between ER+ and ER-subgroups (Fig. 3b) (Additional file 9). These transcriptome changes were validated using RNA-Seq data from the knockdown of *ESR1* and control in MCF7 cells [24] (Fig. S4a in Additional file 2) (Methods). We derived a predictive model with 81 discriminant transcripts that separated ER+ and ER- samples with an average AUC of 0.999 (Fig. 3c). Among the largest PSI changes we found an isoform of the MAP kinase *MAP3K7,* whose long isoform was linked before to apoptosis [25], which we found to be less abundant in ER- samples (Fig. S4b in Additional file 2). Notably, 47 (58%) of the genes with transcripts in this predictive model show differential expression, suggesting a link between estrogen receptor expression and the differential use of transcript isoforms.

**Figure 3.**
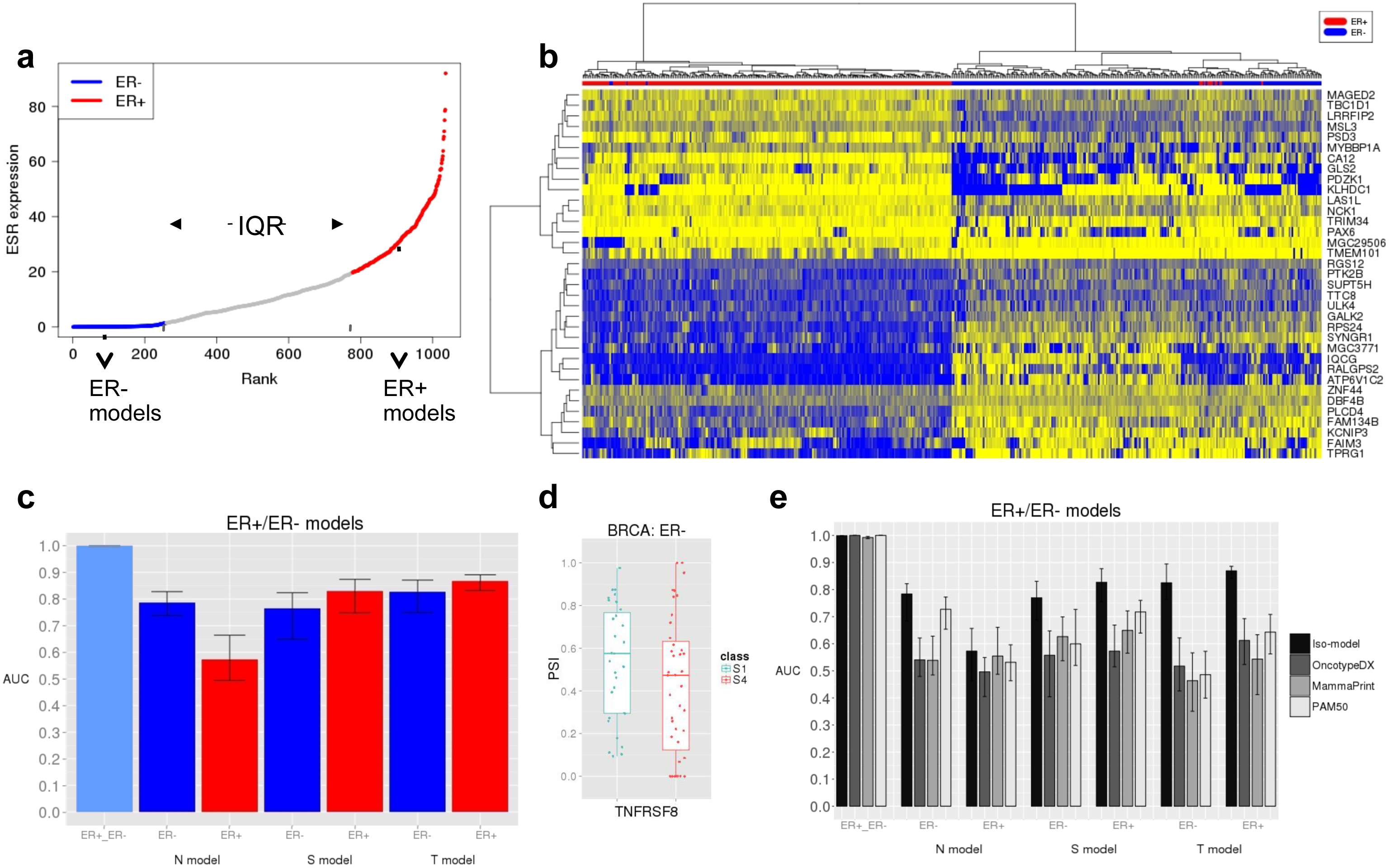
**(a)** Ranking (x axis) of breast tumor (BRCA) samples according to *ESR1* expression (gene TPM) (y axis). ER+ and ER- subsets were defined as the top and bottom 25% of the ranking, respectively, leaving out samples in the inter-quartile range (TQR). **(b)** Heatmap of PSI values, from 0 (blue) to 1 (yellow), for the top 35 isoforms that separate ER+ and ER- subsets. Tsoforms are labeled by gene name (y axis). Samples are clustered according to the PSI values using Euclidean distance and Ward’s method. **(c)** Accuracies in terms of AUC values (y axis) from a 10-fold cross-validation for the transcript isoform signatures for the comparison of ER+ and ER- samples, and for the comparison of early and late N, S and T stages within ER+ or ER- subsets. The variation on each bar indicates the minimum and maximum AUC values. **(d)** PSI distribution of the isoform in *TNFRS8* that changes between early and late S stage in ER- samples (Wilcoxon test p-value = 0.1046). **(e)** Accuracies in terms of AUC values (y axis) from a 10-fold cross-validation for the transcript isoform signatures (Tso-model) and the gene expression signatures OncotypeDX, MammaPrint, and PAM50, indicated in gray scale. Each signature was tested to predict the separation of ER+ and ER- breast tumor samples, or the separation between early and late (N, S and T) stage in ER+ or ER- separately. The variation on each bar indicates the minimum and maximum AUC values.

The observed transcriptome differences between ER+ and ER- subtypes warrant a separation of these two sets to build transcript signatures of stage. Accordingly, we considered early and late stage patients in each ER group separately (Table 3). Since ER- samples show significant differences in survival between early and late stages for N (p-value = 0.005) and S (p-value = 0. 041) annotations (Figs. S4c and S4d in Additional file 2), we expect that signature for stage may be relevant for prognosis. In contrast, ER+ samples do not show any significant differences in survival. Using our feature selection pipeline, we obtained 456 and 249 isoforms that best discriminate between early and late stages in the ER- and ER+ subsets, respectively (Additional file 9). The isoforms for ER- show enrichment in various cancer hallmarks, including DNA repair, Apoptosis and Epithelial-Mesenchymal transition (Fig. S4e in Additional file 2). In contrast, there were no enriched hallmarks associated to the isoforms in the ER+ subset. Building stage signatures as before for ER+ and ER- independently (Additional file 9), we obtained average accuracies of AUC = 0.794 (ER-) and AUC = 0.756 (ER+) (Fig. 3c), with similar values for the precision-recall (Fig. S4f in Additional file 2). Notably, none of the derived signatures showed differential expression at the gene level. Additionally, ER-S-model includes *TNFRS8* (Fig. 3d), a member of the tumor necrosis factor receptor superfamily. Another member of this family, *TNFRSF17* was related before to prognosis in ER- samples [3]. Unlike for the previous models, there were not enough unlabeled samples to perform a blind test. Taken together, these results show that transcript variants can be informative for stage and prognosis in ER negative tumors.

**Table 3.**
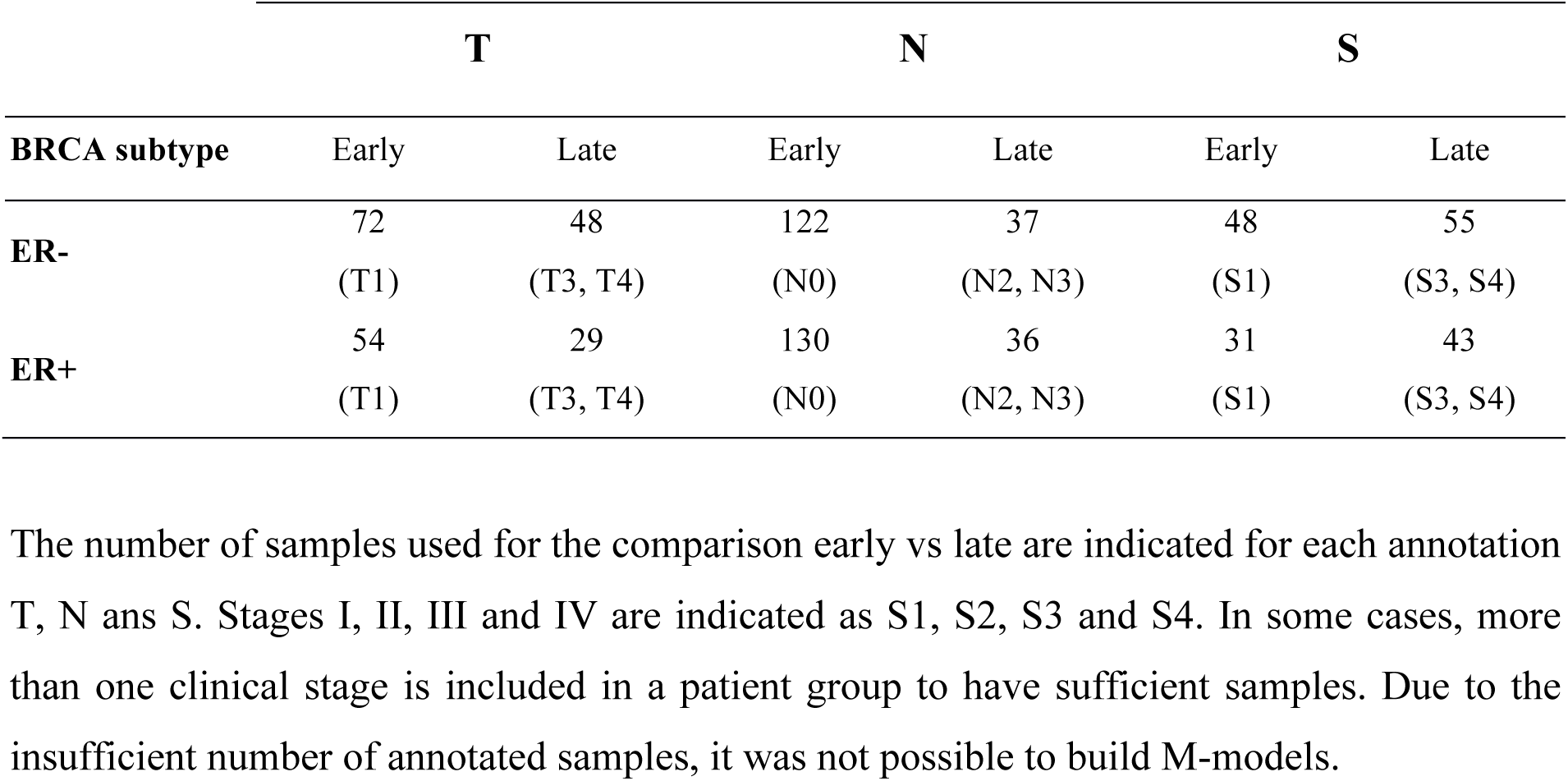
ER-negative (ER-) and ER-positive (ER+) breast tumor subgroups.

|  | **T** |  | **N** |  | **S** |  |
| --- | --- | --- | --- | --- | --- | --- |
| **BRCA subtype** | Early | Late | Early | Late | Early | Late |
| **ER-** | 72<br>(T1) | 48<br>(T3, T4) | 122<br>(N0) | 37<br>(N2, N3) | 48<br>(S1) | 55<br>(S3, S4) |
| **ER+** | 54<br>(T1) | 29<br>(T3, T4) | 130<br>(N0) | 36<br>(N2, N3) | 31<br>(S1) | 43<br>(S3, S4) |

The number of samples used for the comparison early vs late are indicated for each annotation T, N and S. Stages I, II, III and IV are indicated as S1, S2, S3 and S4. In some cases, more than one clinical stage is included in a patient group to have sufficient samples. Due to the insufficient number of annotated samples, it was not possible to build M-models.

We further compared our transcript signatures with known gene expression signatures for breast tumors: OncotypeDX [26], MammaPrint [27], and PAM50 [28] (Methods). Although these signatures were not originally designed to identify tumor stage, they bear predictive value for this purpose [7]. Their accuracies to separate ER+ and ER- subgroups were very similar to our transcript signatures (Fig. 3e). This is expected for PAM50 and OncotypeDX, as they include *ESR1*. We then tested how well the gene signatures differentiate stage within each subset, ER+ or ER-, independently. In general, PAM50 performed better than the two other signatures, except for S in ER- and for N in ER+, where MammaPrint performs better, and for T in ER-, where OncotypeDX performs better (Fig. 3e). Notably, in all cases the transcript signature had better accuracies. We conclude that transcript isoform models can provide relevant information to determine stage and hence complement current clinical signatures.

### Transcript relative abundances characterize an invasive phenotype and survival in melanoma

Clinical outcome of skin cutaneous melanoma (SKCM) remains poor due to its high degree of heterogeneity [29]. The microphthalmia-associated transcription factor *(MITF)* presents highly dynamic expression patterns in connection to proliferation and invasion in melanoma, with relevance for prognosis and therapy [30, 31]. Overexpression and downregulation of *MITF* have been connected to proliferative and invasive phenotypes, respectively [32]. We thus tested whether there are specific transcript signatures linked to these phenotypes that may be predictive of survival. We pooled the top and bottom 25% of melanoma samples according to *MITF* expression into the MITF+ and MITF- sets, respectively (96 samples per set) (Fig. 4a). Although these subsets do not show a significant difference in survival, samples in the top and bottom 10% of *MITF* expression (36 samples per set) show a significant difference, with *MITF* overexpressed samples showing worse prognosis (p=0.029) (Fig. 4b). Our feature selection strategy (Fig. 1a) yielded 2387 discriminant isoforms between MITF+ and MITF-(Fig. 4c) (Additional file 9). We validated these isoforms by comparing their ΔPSI values with those obtained from the knockdown of *MITF* in melanoma cells compared to controls [13] (Fig. S4a in Additional file 2) (Methods). The found discriminant isoforms are enriched for multiple cancer hallmarks, including EMT and the mTOR pathway (Fig. S4b in Additional file 2). To further characterize their differences, we built a predictive model to separate MITF+ and MITF- samples with 72 isoforms, which showed a mean AUC of 0.996 (Methods). This model included a transcript isoform for the cancer *TPM1*, which is highly included in MITF+ and was linked before to tumor growth [33] (Fig. S4c in Additional file 2), as well as for *RAB27A,* a component of the melanosome that is transcriptionally regulated by *MITF* [34] and that is lowly included in MITF+ samples (Fig. S4d in Additional file 2). From this signature, 37 (66%) of the genes involved showed differential expression between MITF+ and MITF- subgroups, pointing to a link between *MITF* expression and differential usage of transcript isoforms in multiple genes.

**Figure 4.**
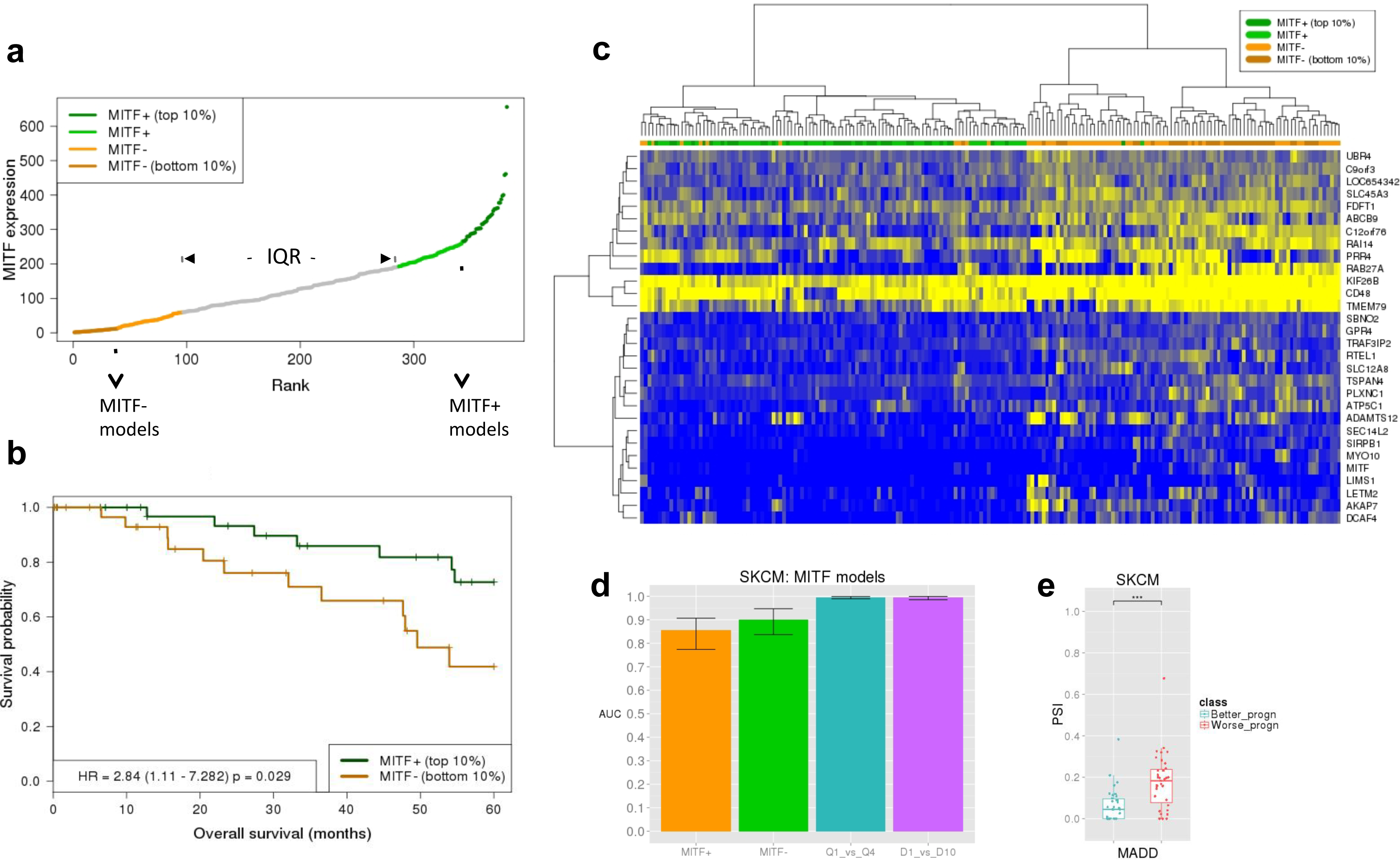
**(a)** Ranking (x axis) of melanoma (SKCM) samples according to *MITF* expression (gene TPM) (y axis). We indicate the top and bottom 10% and 25% of the samples used for analyses. **(b)** Survival (Kaplan-Meyer) plot for the top and bottom 10% of the samples according to the ranking of *MITF* expression. The p-value corresponds to the Cox regression between the two groups and HR indicates the hazards ratio. **(c)** Heatmap of PSI values, from 0 (blue) to 1 (yellow), for the top 30 discriminant isoforms according to |ΔPSI| value between the MITF+ and MITF- subgroups. Isoforms are labeled by gene name (y axis). Samples are clustered according to the PSI values using Euclidean distance and Ward’s method. **(d)** Accuracy given in terms of the distribution of area under the ROC curve (AUC) values (y axis) from a 10-fold cross-validation for (from left to right in the x axis) the survival model for MITF+, MITF- as well as for the separation between MITF+ and MITF- subgroups using 25% (Q1 vs Q4) or 10% (D1 vs D10) of the top and bottom samples in the ranking of *MITF* expression. The bars show the minimum, mean and maximum AUC values. **(e)** Distribution of PSI values for the isoform in *MADD* that is predictive of prognosis in the MITF- subgroup (Wilcoxon test p-value = 7.781e-05).

To test whether the melanoma phenotypes are associated to different transcript transformations during tumor progression, we studied the MITF+ and MITF- sets independently to derive signatures of survival. We selected samples in the top and bottom 40% according to days of survival (36 samples per group) and used our pipeline to calculate the isoforms that best separate each group within each phenotype. The discriminant isoforms in the invasive phenotype (MITF-) were enriched for multiple cancer hallmarks, whereas in the proliferative phenotype (MITF+) presented enrichment only for activation of *KRAS* signaling, which does not appear in the invasive phenotype (Fig. S4b in Additional file 2). We then built predictive models of survival for each subset independently using LMTs (Additional file 9). Cross-validation yielded for MITF+ (34 isoforms) and MITF- (46 isoforms) accuracies of AUC = 0.854 and 0.896, respectively (Fig. 4d) (Fig. S4e in Additional file 2). Notably, the MITF- model includes a transcript isoform for the MAP Kinase-Activating Death Domain gene *MADD* (Fig. 4e), which does not change expression at the gene level. *MADD* is a cancer driver and it was shown before that expression of isoforms that skip exon 16 has anti-apoptotic effects [35]. Interestingly, the PSI of the *MADD* isoform that skips exon 16 is higher in the group with worse prognosis, suggesting that the anti-apoptotic function of *MADD* is related to worse prognosis in invasive melanoma. Taken together, our results provide evidence of distinct transcript abundance patterns linked to melanoma phenotypes and survival.

## Discussion

We described the first systematic analysis of the predictive potential of transcript relative abundances for stage and clinical outcome in multiple solid tumors. We derived novel molecular signatures for 12 different tumor types that can separate tumors according to clinical stage or metastatic status. Importantly, a blind test on patients with unknown stage or metastatic status can separate patients according to survival. Moreover, transcript isoforms provide better accuracies than local alternative splicing events and can describe more complex changes in exon-intron structures. Although a multi-cancer signature of clinical outcome based on gene expression has been proposed [36], our results argue against a generic transcript-based signature for all tumors types. Rather, transcript isoform changes appear linked to tumor-type specific processes, with several of them related to MYC activity, in concordance with recent findings [37].

We observed a widespread association between transcript isoform changes and expression changes in T and S models across tumor types, and around 60% of the genes with differential expression in all models correspond to KIRC and KIRP, indicating that transcript and gene changes are tightly coupled during progression of these tumors. In contrast, this association is low or absent for most tumor types for M and N models. The blind-test showing that patients with predicted metastasis or late N-stage predicted patients have a worse prognosis, was for tumor types for which none of these models show gene expression differences. The value of the transcript signatures is further highlighted when compared to known and newly derived gene expression signatures, or with mixed models combining gene expression and transcript abundances. These results indicate that transcripts signatures provide information independent from gene expression to describe tumor progression, and especially in metastasis and lymph node invasion.

We also extracted prognostic signatures for specific tumor subtypes in breast cancer and melanoma. We reported many significant transcript isoform changes between breast tumors according to estrogen receptor expression and between melanoma samples according to *MITF* expression. Additionally, we observed a widespread association between transcript isoform and expression changes between in relation to estrogen receptor expression and according to *MITF* expression. An interesting possibility is that the activity of these two transcription factor genes could trigger expression and transcript isoform changes in the same genes in these tumors, pointing to new mechanisms of gene regulation worth investigating further. We further derived transcript signatures of stage independently in each sample subset that involved different genes, thereby highlighting the relevance of determining the transcriptome repertoire in tumor samples to derive accurate molecular signatures of tumor progression.

We observed partial reproducibility of the discriminant isoforms in experiments using cell lines. Transcriptional differences between cell lines and tumor tissues are thought to stem from the loss of the stromal and immune components by cells in culture [38]. Our analyses discard an association between the transcript signatures and the composition of stromal and immune cells in the tissue samples. It could be possible that part of the signatures reflect the interaction of tumor cells with their environment in tissues samples, which would be undetectable in cell lines. Our results support the notion that phenotypic states of tumor cells, like invasiveness, may be reflected on the relative abundance of transcript isoforms, may be partly triggered by external cues, such as inflammation or metabolic stress [39]. On the other hand, the observed commonalities between tumor cells and tissues suggest that some of these alterations could be investigated further using cell lines.

It remains to be tested the clinical validity of our findings. Although we have shown that predicted late stages may be associated with worse prognosis and cross-fold validation shows better accuracies in general for transcript signatures, it is not conclusive whether the proposed molecular signatures would actually improve current methodologies of stage determination. Our results indicate that isoform-based M and N models are generally accurate and often better than using gene expression. Those models may be especially useful, as they would indicate a metastasis or lymph node invasion before it is visible by other means. To test this, more validations on independent cohorts would be necessary. However, further studies are currently hampered by the scarcity of large enough datasets with clinical annotation comparable to TCGA [40]. Moreover, the current accuracies of the transcript models may require a larger number of samples to perform prospective studies. Nonetheless, we anticipate that transcript isoforms will be relevant to understand the progression of tumors beyond DNA and gene expression alterations and represent useful novel targets to predict stage and clinical outcome, thereby complementing current molecular approaches in precision cancer medicine.

## Methods

### Datasets

Processed RNA sequencing data from The Cancer Genome Atlas (TCGA) (https://tcga-data.nci.nih.gov/tcga/) was compiled for 12 different tumor types: breast carcinoma (BRCA), colon adenocarcinoma (COAD), head and neck squamous cell carcinoma (HNSC), kidney chromophobe carcinoma (KICH), kidney renal clear cell carcinoma (KIRC), Kidney renal papillary carcinoma (KIRP), lung adenocarcinoma (LUAD), lung squamous cell carcinoma (LUSC), prostate adenocarcinoma (PRAD), skin cutaneous melanoma (SKCM), thyroid carcinoma (THCA) and ovarian carcinoma (OV). The abundance of every transcript per sample was calculated in transcripts per million (TPM) from the transcript-estimated reads counts and the isoform lengths. Genes were defined to be a set of transcripts that overlap in the same genomic locus and strand and share at least one splice-site (Additional file 1). A gene TPM was defined as the sum of TPMs for all transcripts in the gene. The relative abundance of each isoform (PSI), was calculated by normalizing the isoform TPM to the gene TPM. Only genes with a minimum TPM of 0.1 were considered. Additionally, we used RNA-Seq data from the knockdown of *ESR1* and controls in MCF7 cells (GSE53533) [24], from metastatic melanoma cells (SKMel147) and melanocytes (GSE68221) [12], and from the knockdown of *MITF* and controls using non-metastatic melanoma cells (Mel505) (GSE61967) [13]. For each sample, transcript abundances were calculated with Sailfish [41]. Relative abundances (PSI) of transcripts were calculated as above and the ΔPSI values between conditions were calculated as the difference between conditions of the mean values from the replicates. Alternative splicing events and their PSI values were obtained from [42].

### Clinical data

Clinical stage and survival information for patients was obtained from TGCA. We used the available annotation for the TNM staging system (www.cancerstaging.org/), where T followed by a number (1–4) describes the size of the tumor; N followed by a number (1–3) describes spread to lymph nodes according to number and distance; and M followed by 1 or 0 indicates whether the tumor has metastasized or not, respectively. We also considered the numbered stage annotation (S), which goes from 0 to 4, with each number corresponding approximately to a combination of the TNM numbers. When any of the stages were subdivided, only the label of the common class was included (e.g. T1a, T1b and T1c were considered as T1). Only patients with defined stage were used to build the predictive models.

### Selection of relevant features

Only isoforms and events with a difference in mean relative abundance (PSI) of at least 0.1 in absolute value between the compared patient subgroups were considered to calculate discriminant isoforms. To obtain discriminant genes, those with log-fold change of the mean gene TPM values between the two groups greater than 2 were considered. Next, a subsampling approach was used to compare two patient groups through 100 iterations, by extracting the same number of samples from each group randomly from the input dataset, using a minimum of 10 samples per group. For pooled tumor types, the same number of samples per tumor type was selected at each iteration step. At each iteration step, three different univariate discriminant measures were applied (see below), and a permutation of the group labels was performed and the univariate measures re-calculated. After 100 iterations, and for each univariate measure, two distributions of 100 points each are produced for each transcript, corresponding to the observed and expected values. Transcripts with a positive difference of the means of the two distributions for all three measures were considered discriminant and were kept for further analysis.

We applied the following information-based measures in the subsampling: information gain (*IG*), gain ratio (*GR*) and symmetrical uncertainty (*SU*). *IG* is defined as the mutual information between the group labels of the training set *S* and the values of a feature (or attribute) A, e.g. an isoform: *IG*(*S*, *A*) = *MI*(*S*,*A*) = *H*(*S*) - *H*(*S|A*), where *H*(*S*) is Shannon’s entropy according to the two sample classes, and *H*(*S|A*) is the conditional entropy of *S* with respect to the attribute A. *GR* is the mutual information of the group labels and the attribute, normalized by the entropy contribution from the proportions of the samples according to the partitioning by the attribute: *GR*(*S*,*A*) = *MI*(*S*,*A*) / *H*(*A*). Finally, *SU* provides a symmetric measurement of feature correlation with the labels and it compensates possible biases from the other two measures: *SU*(*S*,*A*) = *2MI*(*S*,*A*) / (*H*(*S*) + *H*(*A*)) [43]. The group labels are the clinical stages (early, late), survival groups (low, high), or phenotype group (invasive, proliferative); and the attribute values are the PSI values for transcript isoforms or alternative splicing events, or the gene TPM values for gene expression analyses. The continuous PSI or TPM values were discretized as previously described [44].

### Cancer hallmarks and drivers

Enrichment analysis of the 50 cancer hallmarks from the Molecular Signatures Database v4.0 [45] was performed with the discriminant isoforms. For each hallmark, a Fisher exact test was performed with the genes with selected isoforms using as controls genes expressed (TPM>0.1) and with multiple transcripts A Benjamini-Hochberg correction was applied and only cases with FDR < 0.05 were kept. Known and predicted cancer drivers were obtained as described in [42].

### Predictive signatures

Transcript isoforms that showed a positive difference between the means of the 100 observed and the 100 randomized values for all three univariate measures (*IG*, *GR*, *SU*) were analyzed with a Correlation Feature Selection (*CFS*) (Hall 2000). This selects transcripts with similar discriminating power but lower redundancy among them (Hall 2000), thereby mitigating the problem of overfitting. This was repeated for each comparison between clinical stages, survival groups, or tumor subtypes. Using the selected transcript isoforms, a Logistic Model Tree (*LMT*) was built with Rweka [46]. *LMTs* are classification trees with logistic regression functions at the leaves. The accuracy of the classifiers was evaluated using the area under the receiver-operating characteristic (ROC) curve or AUC. Additionally, we considered the area under the precision-recall curve (PRC). AUC and PRC take values between 0 (worst prediction) and 1 (best prediction). These values were estimated for each classifier through a 10-fold cross validation, repeated 100 times. The same approach was used for gene, event, and mixed models. To apply known gene expression signatures to our sample groups we used robust Z-scores per gene and per sample as described before [42]. These values were the used for the genes in various signatures [21,26,27,28]. As before, accuracies were estimated using a 10-fold cross-validation to calculate AUC and PRC values.

### Blind tests

For samples without stage annotation, which were not used to build the models, we predicted the missing stage (early/late) or metastatic state, using the corresponding model for the same tumor type. These newly predicted samples were then aggregated per clinical class according early and late, or metastatic and non-metastatic, to test the survival differences between groups. The blind test was performed using only those tumor types that already showed significant differences in the survival between early and late stages for the annotated samples (Table 2). This analysis was not performed for T-models, as all samples had a T annotation.

### Differential expression analysis

We performed differential expression (DE) analysis for all genes between the different groups considered in this analysis, using the same method as described previously [41]. Genes were considered differentially expressed if the absolute value of the log2-fold change was greater than 0.5 and corrected p-value < 0.05. Results can be found in Additional_files_6 and 10.

### Survival analysis

Survival curves were calculated with the Kaplan-Meier method and compared between patient subsets using a Cox proportional hazards regression model [47]. Survival was measured as date of death minus collection date for deceased patients and as last contact date minus collection date for the other patients.

### Stromal and Immune cell content analysis

To estimate a stromal and immune signature for a set of samples from a tumor type, we collected a list of stromal and immune signature genes based on [18]. We transformed the RSEM read counts of these two gene lists into a gene set score using GSVA [48] for each sample. Using the resulting scores per sample, we then calculated the Pearson correlations of the stromal and immune GSVA scores with the transcript isoform PSIs using all tumor samples, including intermediate stages.

## List of abbreviations used

PSI: percent/proportion spliced in, IG: information gain, GR: gain ration, SU: symmetrical uncertainty, CFS: correlation feature selection, ER: estrogen receptor, LMT: logistic model tree, ROC: receiver operating characteristic, AUC: area under de ROC curve, PRC: area under the precision-recall curve.

## Authors’ contributions

EE proposed and supervised the study, JLT carried out work. AP and ES contributed with some of the software components and processed datasets. JLT and EE wrote the paper with inputs from ES. All authors read and approved the manuscript.

## Description of additional data files

The following additional data are available with the online version of this paper:

Additional file 1: Patient data, blind predictions and gene-isoform annotations.

Additional file 2: Figures S1, S2, S3, S4 and S5.

Additional file 3: Discriminant transcript isoforms between stages and patient groups. Additional file 4: Enriched cancer hallmarks.

Additional file 5:Transcript signatures of tumor stage

Additional file 6: Correlation of predictive isoforms with stromal and immune scores. Additional file 7: Transcript signatures in pooled tumor types.

Additional file 8: Predictive signatures based on gene expression and splicing events. Additional file 9: Models in ER+/ER- breast and MITF+/MITF- melanoma tumors.

## Acknowledgements

We would like to thank Victor Moreno, Jun Yokota, and Rubén Pio, Angel Rubio and Luis Montuenga for useful discussions and comments on earlier versions of the manuscript. This work was supported by grants BIO2014-52566-R and Consolider RNAREG (CSD2009-00080) from the MINECO (Spanish Government), by AGAUR (2014-SGR1121) and by the Sandra Ibarra Foundation for Cancer (FSI2013).

## Figure Legends

**Figure S1**. **(a)** Information-based feature selection methods provide a robust and conservative measure of the discriminant power of features. Left panels show plots comparing the information gain (IG) (upper panel), gain ratio (GR) (middle panel) and symmetrical uncertainty (SU) (lower panel) (y axes) with the Wilcoxon-test p-value after multiple-testing correction using Benjamini-Hochberg method (x axes) for the distribution of PSI values for two patient subgroups. In this case, the data corresponds to the comparison between ER+ and ER- breast tumor samples, subsampling 20 patients per group. Each dot corresponds to one isoform in each of the subsamples. Right panels: in red we show the distributions of IG (upper panel), GR (middle panel) and SU (lower panel) values for the comparison of a transcript from MAP3K7 between ER+ and ER- samples with using 100 subsamples of size 20. In blue we show the same values in the comparison between the groups after shuffling the labels. **(b)** Enriched hallmarks (y axis) for each stage class, metastasis (M), tumor size (T), lymph-node involvement (N) and overall staging (S), in each tumor type (x axis), using all discriminant isoforms found in each case. Only significant cases (corrected Fisher test p-value < 0.05) are shown. **(c)** Comparison of the ΔPSI values for the discriminant isoforms between metastatic and non-metastatic SKMC samples (x axis) with the ΔPSI values from the comparison of the metastatic (SKMel147) and non-metastatic (Mel505) melanoma cells (y axis). We indicate in blue or red those isoforms with the same or opposite change direction, respectively. Dark and light colors indicate |ΔPSI|>0.1 and |ΔPSI|<0.1, respectively. The correlation (Pearson R) is given for isoforms in dark blue. **(d)** Accuracy of the models in terms of the areas under the precision-recall curves (PRC) (y axis) for the late-stage classes (i.e. precision is measured as the proportion of predicted late stage samples that are correctly predicted). The bars show the minimum, mean and maximum values of the area of the precision-recall curves. Some models are absent due to lack of sufficient samples (Table 1).

**Figure S2**. **(a)** PSI distributions of some of the transcript isoforms in the derived predictive signatures. From left to right, *PAX6* isoform in the KIRP T-model for KIRP (Wilcoxon test p-value = 2.695e-06), *MKNK1* isoform in the KIRP N-model (Wilcoxon test p-value = 0.0004), *TM6SF1* isoform in the SKCM M-model (Wilcoxon test p-value = 1.813e-05), *PRDM16* isoform (Wilcoxon test p-value = 0.0001) and *PTKB* isoform (Wilcoxon test p-value = 0.005) in BRCA S-model. The y-axis indicates the PSI value in each sample separated according to early and late stages (x-axis). **(b)** Left panel: XY-plot of the PSI values (y axis) of the *ENAH* isoform that appears in the T-models of KIRP and COAD, and the stromal score (x axis), across all COAD tumor samples. Pearson correlation with stromal score R=-0.59 and with immune score R=-0.41. Right panel: PSI distribution of the same *ENAH* isoform in early and late T-stages in KIRP and COAD (Wilcoxon test p-value < 0.001).

**Figure S3**. **(a)** Accuracies of the transcript isoform models (T) compared to the gene (G) and mixed (M) combining isoform and gene information. Accuracies are given as boxplots for the distribution of AUC values (y axis) from a 10-fold cross-validation for each tumor type (x axis) for the metastasis (M), overall stage (S), lymph node invasion (N) and tumor size (T) models (panels from upper left to lower right). Tumors for which stage data was missing are not shown (see Table 1 of the manuscript). **(b)** and **(c)**: Survival (Kaplan-Meyer) plot associated to the blind test for N and S stage, respectively using the mixed model combining isoform and gene information. The plot indicates the survival percentage (y axis) versus survival in months (x axis) based on the predicted stage on the unannotated samples using the classifier for each corresponding tumor type. The p-value in each plot corresponds to the Cox regression between the two groups and HR indicates the hazards ratio

**Figure S4**. **(a)** Comparison of the ΔPSI values for the discriminant isoforms between ER+ and ER- samples (x axis) with the ΔPSI values from the comparison of the control and knockdown of *ESR1* in MCF7 cells (y axis). We indicate in blue or red those isoforms with the same or opposite change direction, respectively. Dark and light colors indicate |ΔPSI|>0.1 and |ΔPSI|<0.1, respectively. The correlation (Pearson R) is given for isoforms in dark blue. From the 2337 transcript isoforms with expression in the MCF7 experiments, 1123 (48%) show PSI changes in the same direction and 328 of them with |ΔPSI| > 0.1. **(b)** PSI distribution of the *MAP3K7* isoform that changes significantly between the ER+ and ER- BRCA sets (Wilcoxon test p-value < 2.2E-16). Plots **(c)** and **(d)** show the survival (Kaplan-Meyer) curves for the ER- samples according to early and late N and S stages, respectively. The p-value in each plot corresponds to the Cox regression between the two groups and HR indicates the hazards ratio. **(e)** Enriched cancer hallmarks for the set of discriminant isoforms between ER+ and ER- subsets (ER+_ER-) and for the set of isoforms separating early and late stages in ER- (ER-). For this latter comparison isoforms associated to N, T and S stages were combined into early and late subgroups. **(f)** Accuracy of the models in terms of the areas of the precision-recall curves (PRC) (y axis) for the comparison between ER+ and ER- subgroups and for the comparison of early vs late stage classes in each subtype, ER+ and ER-, for N, S and T annotation. The precision is measured as the proportion of predicted late stage samples or ER+ samples that are correctly predicted. The bars show the minimum, mean and maximum values of the area of the precision-recall curves.

**figure S5**. **(a)** Comparison of the ΔPSI values for the discriminant isoforms between MITF+ and MITF- melanoma tissue samples (x axis) with the ΔPSIs obtained from the comparison of the control and knockdown of *MITF* in Mel505 cells (y axis). We indicate in blue or red those isoforms with the same or opposite change direction, respectively. Dark and light colors indicate |ΔPSI|>0.1 and |ΔPSI|<0.1, respectively. The correlation (Pearson R) is given for isoforms in dark blue. From the total of 2279 discriminant isoforms for which we found expression in the cell lines, 1050 (46%) show a ΔPSI change in the same direction, with 865 of them having |ΔPSI|>0.1. **(b)** Enriched cancer hallmarks (y axis) (corrected Fisher test p-value < 0.05) using the discriminant isoforms in the comparison MITF- vs MITF+ and comparing low and high survival subgroup of patients within each subtype MITF+ or MITF-. Enriched hallmarks were the same using the top and bottom 10% or 25% samples according to *MITF* expression to define the subtypes. **(c)** PSI distributions of the *TPM1* isoform (left panel) and *RAB27A* isoform (right panel) that separate the two melanoma subtypes, MITF+ and MITF- (Wilcoxon test p-values = 5.293e-9 and 4.86e-12, respectively). The plots indicate the PSI values (y-axis) for the isoforms in MITF+ and MITF- samples (x-axis). **(d)** Genomic locus for RAB27A indicating the annotated isoforms; uc002acr.2 decreases PSI in MITF+ (Fig. S4c in Additional file 12), whereas uc002acp.2 increases PSI in MITF+ (Additional file 10). **(e)** Accuracy given in terms of the areas under the precision-recall curves (PRC) (y axis) from a 10-fold cross-validation for (from left to right in the x axis) the survival model for MITF+, MITF- as well as for the separation between MITF+ and MITF-subgroups using 25% (Q1 vs Q4) or 10% (D1 vs D10) of the top and bottom samples in the ranking of *MITF* expression The bars show the minimum, mean and maximum values of the area of the precision-recall curves.

